## Supplementary Figure 1 for "Age-related changes in Tau and Autophagy in human brain in the absence of neurodegeneration"

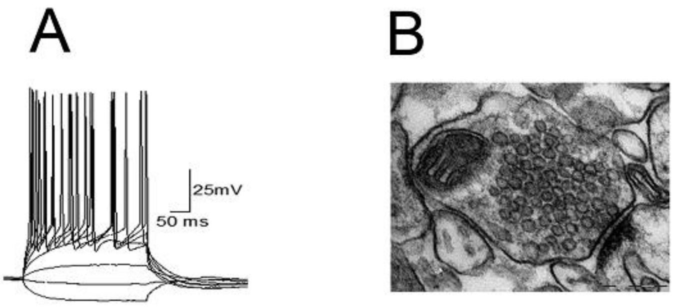


**Suppl Figure 1. Electrophysiological recordings and ultrastructure of cortical resected tissue.** Whole-cell patch clamp recording from a layer II-III human cortical pyramidal neuron showing active voltage responses to current injection (A). Electron micrograph showing a cortical synapse Scale bar 200 nm (B).
