## Supplementary Figure 2 for "Age-related changes in Tau and Autophagy in human brain in the absence of neurodegeneration"

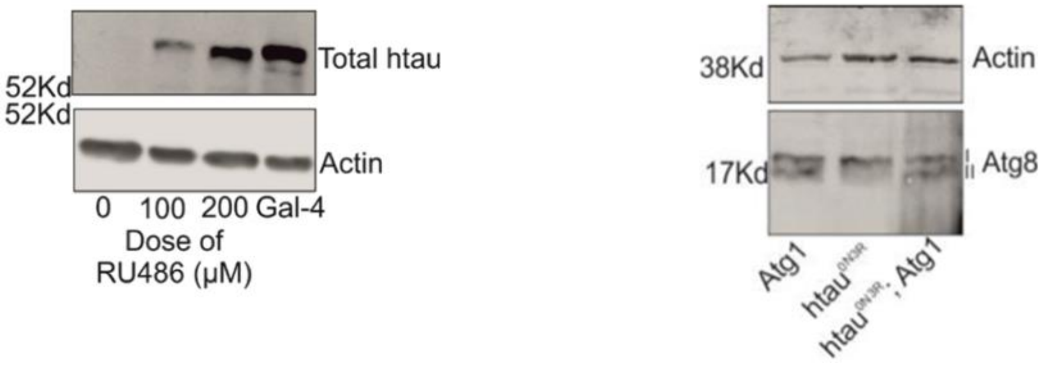


**Supplementary Figure 2: Unexpressed controls. (**A) *Elav*-GeneSwitch (GS) expression system allows for temporal control of pan-neural hTau$^{0N3R}$ expression following introduction of RU486 upon eclosion. No expression is evident when no drug is given (0μM RU486) and a dose dependent increase in tau expression is seen with increasing doses of RU486. 200μM RU486 gives tau expression that is comparable to that seen with the standard *Elav*-GAL4 so this dose was chosen for all the experiments in this study. (B) Atg8 staining in *Elav*-GS/*UAS*-Atg1, *Elav*-GS/hTau$^{0N3R}$ and *Elav*-GS/ hTau$^{0N3R}$; Atg1 transgenics after treatment with 200μM RU486 upon eclosion. During autophagy, Atg8-I is conjugated with a lipid moiety and the lipidated form (Atg8-II) is recruited to autophagosomal membranes. An elevated Atg8-II level therefore serves as an indicator of autophagy activation. Atg8-II levels are evident in Atg-I and hTau$^{0N3R}$;Atg-1 flies but not to any great extent in hTau$^{0N3R}$ alone flies.
