## Supplementary Figure 3 for "Age-related changes in Tau and Autophagy in human brain in the absence of neurodegeneration"

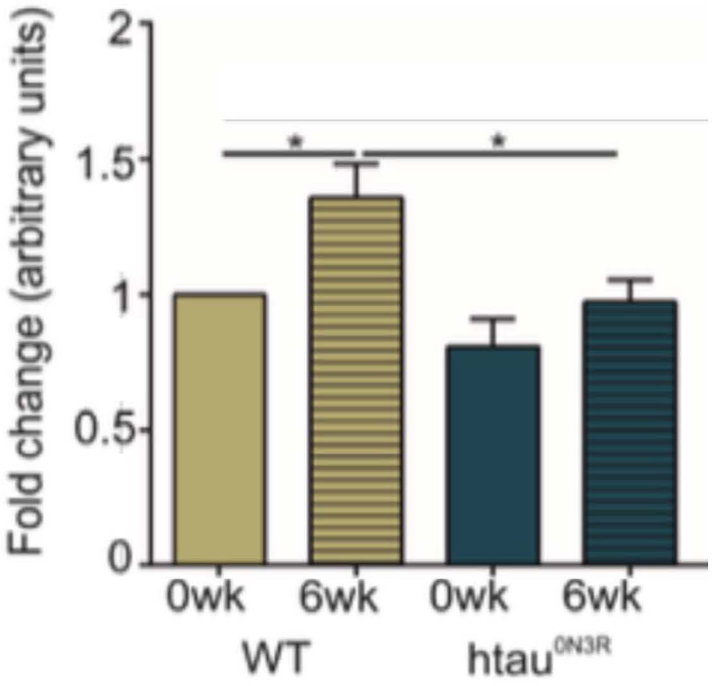


**Supplementary Figure 3: Impact of hTau expression on endogenous dTau.** *Drosophila* tau (dTau) was probed using an antibody specific only to dTau (gift from Johnston lab University of Cambridge) and in wild type (wt) OreR flies, dTau levels were found to increase with age ($p=0.0193$). This age-related increase was not evident upon Elav-Gal4 expression of $hTau^{0N3R}$. The dTau expression in 6wk wt flies that are expressing $hTau^{0N3R}$ was significantly less than that evident in 6wk wt flies ($p=0.0309$) (data is fold change compared to dTau levels in wt flies at 0wk; unpaired two tailed t-test; n=7-14)
