## Supplementary Figure 4 for "Age-related changes in Tau and Autophagy in human brain in the absence of neurodegeneration"

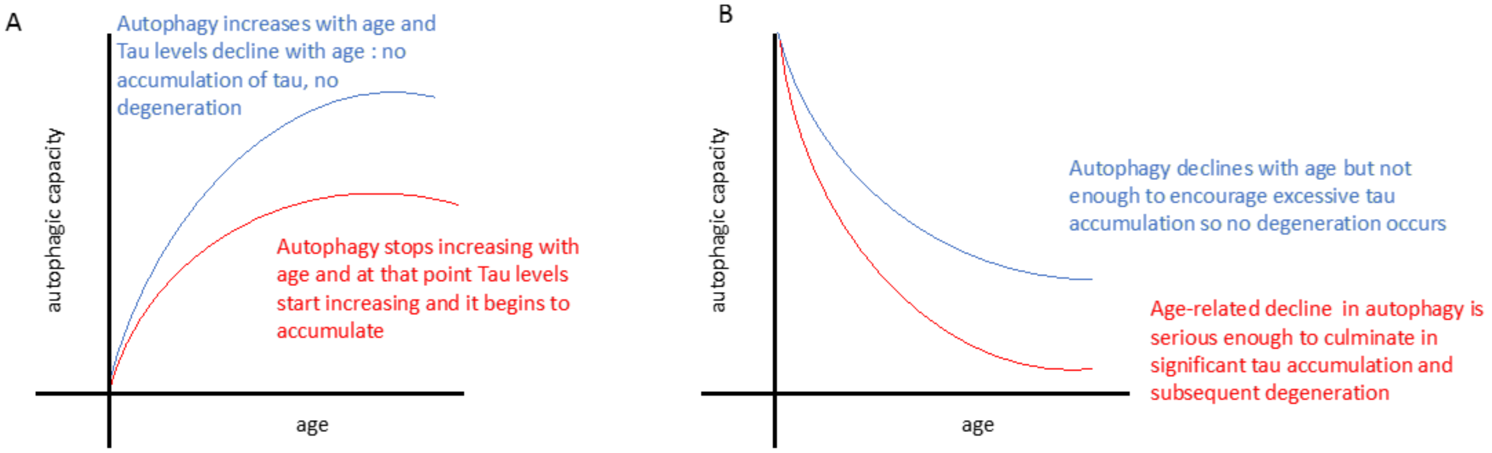


**Supplementary Figure 4. Two hypotheses for how autophagy may change with age and influence accumulation of misfolded proteins:** (A) In individuals with a healthy autophagic capacity, autophagic responses increase with age to deal with age-related insults. This may contribute to a decline in soluble phosphorylated tau levels and protect from tau accumulation and subsequent degeneration in healthy long-lived elderly subjects who do not develop tauopathy (blue line). In those elderly where an age-related autophagic response to aging insults begins to decline, misfolded proteins like tau start to accumulate and lead to degeneration and these individuals develop tauopathy (red line). (B) An age-related decline in autophagic capacity is evident in all elderly but in some subjects (blue line) this is not significant enough to encourage tau accumulation and they do no develop degeneration or tauopathy; in other subjects (red line), the age-related impairment occurs to a greater extent and this precipitates accumulation of misfolded proteins and pathogenesis of neurodegenerative diseases like tauopathy.
